# A New Paradigm for Investigating Real-World Social Behavior and its Neural Underpinnings

**DOI:** 10.1101/2021.12.26.474173

**Authors:** Arish Alreja, Michael J. Ward, Qianli Ma, Brian E. Russ, Stephan Bickel, Nelleke C. Van Wouwe, Jorge A. González-Martínez, Joseph S. Neimat, Taylor J. Abel, Anto Bagić, Lisa S. Parker, R. Mark Richardson, Charles E. Schroeder, Louis–Philippe Morency, Avniel Singh Ghuman

**Affiliations:** Center for the Neural Basis of Cognition, Carnegie Mellon University and University of Pittsburgh; Neuroscience Institute, Carnegie Mellon University; Machine Learning Department, Carnegie Mellon University; Department of Neurological Surgery, University of Pittsburgh; David Geffen School of Medicine, University of California Los Angeles; Language Technologies Institute, Carnegie Mellon University; Nathan Kline Institute for Psychiatric Research; The Feinstein Institutes for Medical Research, Dep. of Neurosurgery and Neurology, Northwell Health; Department of Neurological Surgery, University of Louisville; Brain Institute, University of Pittsburgh; Department of Neurology, University of Pittsburgh; School of Public Health, University of Pittsburgh; Department of Neurosurgery, Harvard Medical School and Massachusetts General Hospital; Departments of Neurosurgery and Psychiatry, Columbia University; Departments of Psychology, Neurobiology, and Psychiatry, University of Pittsburgh

## Abstract

Eye tracking and other behavioral measurements collected from patient-participants in their hospital rooms afford a unique opportunity to study immersive natural behavior for basic and clinical translational research. We describe an immersive social and behavioral paradigm implemented in patients undergoing evaluation for surgical treatment of epilepsy, with electrodes implanted in the brain to determine the source of their seizures. Our studies entail collecting eye tracking with other behavioral and psychophysiological measurements from patient-participants during unscripted behavior, including social interactions with clinical staff, friends and family, in the hospital room. This approach affords a unique opportunity to study the neurobiology of natural social behavior, though it requires carefully addressing distinct logistical, technical, and ethical challenges. Collecting neurophysiological data synchronized to behavioral and psychophysiological measures helps us to study the relationship between behavior and physiology. Combining across these rich data sources while participants eat, read, converse with friends and family, etc., enables clinical-translational research aimed at understanding the participants’ disorders and clinician-patient interactions, as well as basic research into natural, real-world behavior. We discuss data acquisition, quality control, annotation, and analysis pipelines that are required for our studies. We also discuss the clinical, logistical, and ethical and privacy considerations critical to working in the hospital setting.

## Introduction

Real-world behaviors such as social interactions are traditionally studied using simplified laboratory conditions in order to control for inherent natural complexities. Real-world environments offer the opportunity to study behavior, and its physiological correlates, in ecologically valid settings. Technological advances in recent decades have enabled us to capture and analyze critical behavioral and physiological variables in real time, over long periods of time, with greater fidelity than ever before (Jacob Rodrigues et al., 2020; Johnson and Andrews, 1996; Topalovic et al., 2020; Wilhelm et al., 2006) to enable modeling real-world variability and complexity through large datasets using modern computational methodology. Doing so in real-world environments allows us to convert real-world complexities from problems to assets, which can prove transformative for understanding natural behavior and its relationship to physiology (Holleman et al., 2020; Matusz et al., 2019; Powell and Rosenthal, 2017; Zaki and Ochsner, 2009).

The inpatient hospital environment is a distinctive real-world setting for investigating the relationship between behavior and physiology. It features monitoring of physiological data (electrocardiograms, electromyograms, heart rate, blood pressure, neural recordings, etc.) as part of standard care that can be augmented with behavioral monitoring (eye–tracking, egocentric video and audio recording, etc.). It also offers the opportunity to observe the relationship between behavior, perception, and physiology before, during, and after clinical events relevant to the patients’ pathology. ***From a clinical perspective***, a deeper grasp of the relationship between behavior and physiology accompanying clinical events has broad implications for diagnostics and our understanding of physiological-behavioral relationships in clinical disorders (Clark et al., 2019; Vigier et al., 2021; Wolf and Ueda, 2021). In addition, the hospital setting provides the opportunity to capture key caregiver–patient interactions, whose salience for patients in such an environment cannot be overstated (Jhalani et al., 2005; Pickering et al., 2002). Modeling these interactions has deep implications in terms of understanding joint clinical decision-making, clinical information transfer, patient outcomes, patient satisfaction and the informed consent process in ways that cannot be replicated in controlled lab environments (Finset and Mjaaland, 2009; Girard et al., 2021; Kiesler and Auerbach, 2006; Muszynski et al., 2020; Weilenmann et al., 2018). ***From a basic science perspective***, the inpatient hospital environment also offers a compelling immersive environment to advance basic knowledge by studying natural behavior, such as interactions with friends and family, clinicians, eating, reading etc., in patients that have simultaneous behavioral, physiological and psychophysiological monitoring (Hogan and Baucom, 2016).

Real-world behavior encompasses a multitude of physiological and behavioral processes unfolding at different timescales, which are affected by ‘change events’ in the environment itself (Shiffman et al., 2008). This makes them challenging to study. Successfully studying the relationship between behavior and physiology in such settings requires extracting meaningful insights from data that are rich, complex and heterogeneous in nature and varied in time. Inpatient hospital settings are subject to these considerations, as well as the additional complexity of hospital environments where unpredictable and potentially adverse events may unfold for patients. In addition, they give rise to ethical considerations that include patient privacy and well-being (and potentially the privacy of others), and the confidentiality of clinical information and doctor–patient interactions (Roter and Hall, 1989).

This paper presents methodology for collecting behavioral and physiological data in epilepsy patients who undergo extra–operative invasive monitoring for seizure localization. Patients are implanted with intracranial electrodes (superficial, depth or a combination of both) and then are admitted to the Epilepsy Monitoring Unit (EMU) for 1–2 weeks for clinical identification of the epileptogenic zone and for functional mapping. This clinical setting presents a unique opportunity to capture behavioral data (eye–tracking using eye–tracking glasses, audio, and video recordings) synchronized with neural activity recorded by intracranial electrodes implanted in the patient’s brain, during real-world social interactions with friends, family, clinicians and researchers. We discuss the privacy and ethical considerations that arise in this paradigm and how they can be addressed, as well as logistical challenges such as fitting seizure prone patients, who have significant head bandaging protecting their implantation sites, with eye–tracking glasses to collect data in a safe and robust manner. Finally, we describe data preprocessing and data fusion pipelines that can be used to construct a high-quality multimodal data set that blends real-world social behavior and neural activity, allowing us to study the neural correlates of real-world social and affective perception in the human brain.

## Materials & Methods

### Participants

A total of 6 patients (4 men, 2 women) underwent surgical placement of subdural electrocorticographic electrodes (ECoG) or stereoelectroencephalography (SEEG) depth electrodes as standard of care for epileptogenic zone localization. Together ECoG and SEEG are referred to here as iEEG. The ages of the participants ranged from 22 to 64 years old (mean = 37 years, SD = 13.47 years). No ictal events were observed during experimental sessions

### Informed Consent

All participants provided written informed consent in accordance with the University of Pittsburgh Institutional Review Board. The informed consent protocols were developed in consultation with a bioethicist (Dr. Lisa Parker) and approved by the Institutional Review Board of the University of Pittsburgh. Audio and video of personal interactions were recorded during experimental sessions. Our protocol incorporated several measures to ensure privacy considerations and concerns could be addressed based on the preferences of individual participants. First, the timing of recording sessions was chosen based on clinical condition and participant preference, to ensure that they were comfortable with recording of their interactions with the visitors present (and/or expected to be present). Second, all visitors present in the room were notified about the nature of the experiment at the beginning of each recording session and given the opportunity to avoid participation. Third, a notification was posted at the entrance of the patient room informing any entrants that an experiment was being conducted where they might be recorded so that they could avoid entering if they chose to. It is notable that there are no reasonable expectations of privacy other than for the patient, and this work was considered to meet the criteria for waiver of informed consent for everyone other than the participants themselves. Finally, at the end of each experimental recording, participants were polled to confirm their consent with the recording being used for research purposes, and offered the option to have specific portions (e.g., a personal conversation) or the entire recording deleted if they wished. Thus, explicit “ongoing consent” was acquired through written informed consent at the beginning and end of each session; providing participants the opportunity both affirm their willingness to participate and to consider the content of the recordings before giving final consent. None of our participants thus far have asked to have recordings partially or fully deleted after the recording session was complete.

### Electrode Localization

Coregistration of grid electrodes and electrode strips was adapted from the method of Hermes et al. (2010). Electrode contacts were segmented from high-resolution postoperative CT scans of participants coregistered with anatomical MRI scans before neurosurgery and electrode implantation. The Hermes method accounts for shifts in electrode location due to the deformation of the cortex by utilizing reconstructions of the cortical surface with FreeSurfer^TM^ software and co-registering these reconstructions with a high-resolution postoperative CT scan. All electrodes were localized with Brainstorm software (Tadel et al., 2011) using postoperative MRI coregistered with preoperative MRI images.

### Data Acquisition

Multimodal behavioral data (audio, egocentric video, and eye–tracking) as well as neural activity from up to 256 iEEG contacts can be recorded simultaneously during unscripted free viewing sessions in which participants wore eye–tracking glasses while they interacted with friends and family visiting them, clinicians and hospital staff responsible for their care, and members of the research team. In addition, participants also engaged in other activities like eating meals, reading, and watching television. The type and duration of activities varied across different recording sessions. The timing and duration of recording sessions were determined based on clinical condition, participant preference and to coincide with the presence of visitors in the hospital room, where possible.

Behavioral data were captured by fitting each participant with SensoMotoric Instrument’s (SMI) ETG 2 Eye Tracking Glasses (Fig. 1.a,c). An outward facing egocentric camera recorded video of the scene viewed by participants at a resolution of 1280 x 960 pixels at 24 frames per second (Fig. 1.b). Two inward facing eye–tracking cameras recorded eye position at 60 Hz (Fig. 1.c,d). Audio was recorded at 16 KHz (256 Kbps) using a microphone embedded in the glasses. SMI’s iView ETG server application, running on a laptop received and stored streaming data for all three modalities from the eye–tracking glasses by way of a USB2.0 wired connection. The iView ETG software also served as an interface for researchers to calibrate the eye-tracking glasses to each participant with a 3 point calibration procedure that enabled the accurate mapping of eye–tracking data to specific ‘gaze’ locations on video frames, and to initiate and stop the recording of behavioral data.

**Figure 1:**
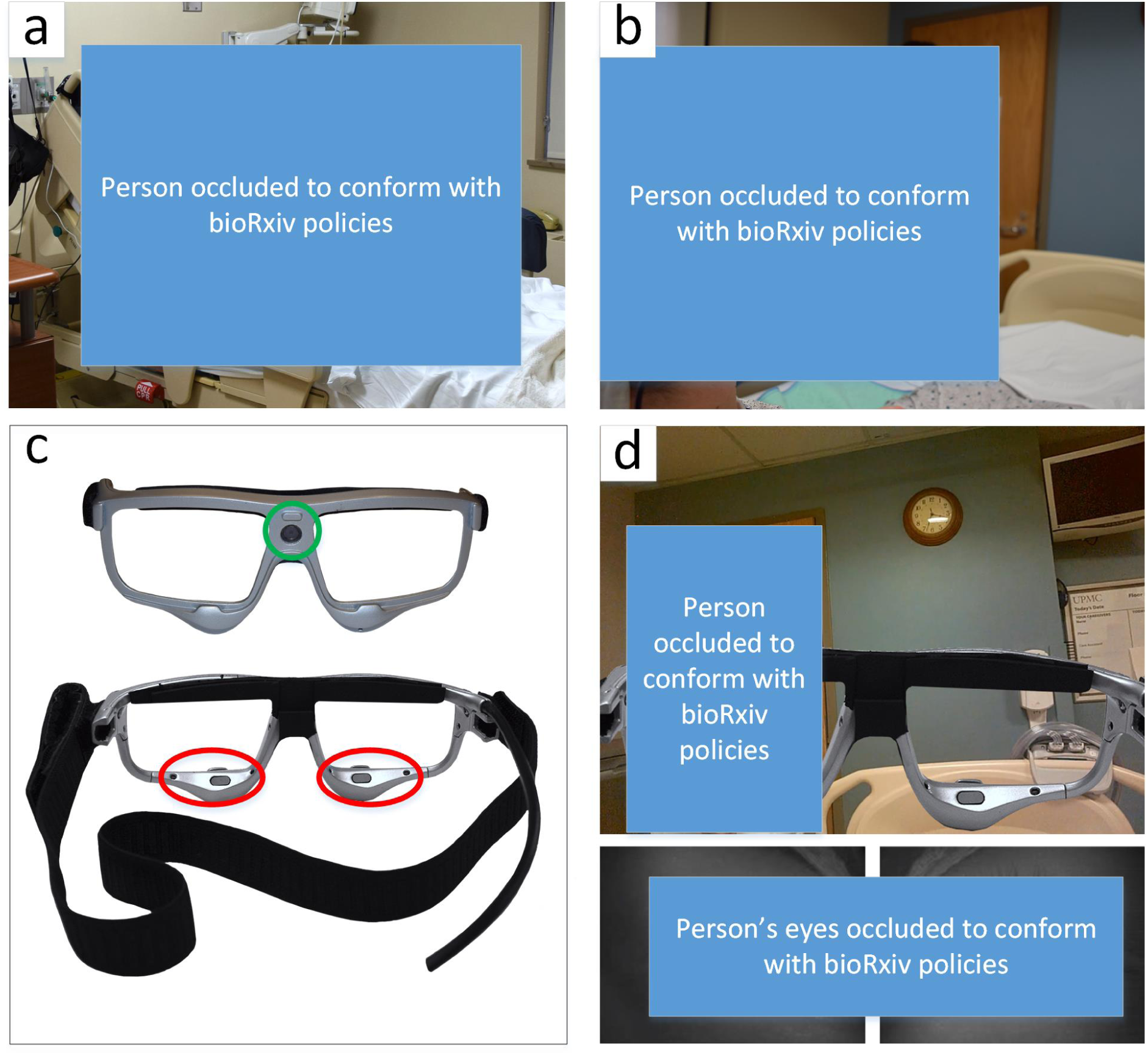
**a)** A participant in the UPMC Epilepsy Monitoring Unit implanted with iEEG electrodes, secured with bandaging, and fitted with with SensoMotoric Instrument’s (SMI) ETG 2 Eye Tracking Glasses that have been modified with an ergonomic Velcro strap. **b)** An over the shoulder view of the participant and the visual scene during an interaction with a researcher. **c)** Front (top) and Back (bottom) view of the SMI ETG 2 Eye Tracking Glasses with the egocentric video camera (green circle) and inward facing eye–tracking cameras (red ellipses). **d)** A snapshot of the participant’s view (top) through the SMI ETG 2 Eye Tracking Glasses corresponding to panel b), and their eye movement (bottom) captured by the inward facing eye–tracking cameras.

Electrophysiological activity (Field Potentials) can be recorded from up to 256 iEEG electrodes at a sampling rate of 1 KHz using a Ripple Neuro’s Grapevine Neural Interface Processor (NIP) (Fig. 2). Common reference and ground electrodes were placed subdurally at a location distant from any recording electrodes, with contacts oriented toward the dura.

**Figure 2:**
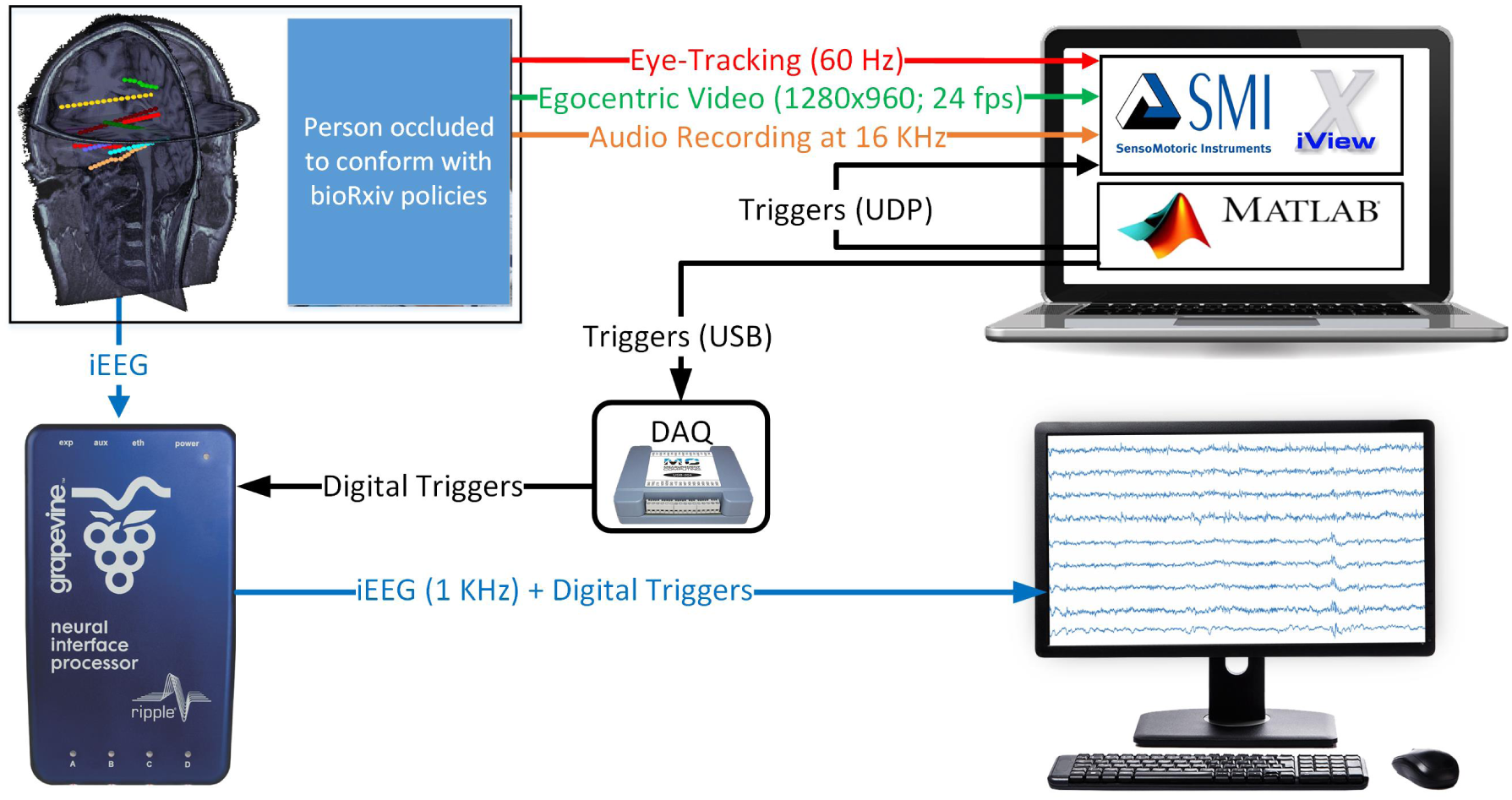
A system diagram of the experimental setup for the collection of synchronized behavioral (egocentric video, eye–tracking and audio) and physiological (iEEG recordings) from participants during real world social interactions. The green, red and blue lines represent egocentric video (1280×960 pixels; 24 fps), eye–tracking (60 Hz), and audio (16 KHz). Digital Triggers, represented by black lines, are inserted in the eye–tracking and iEEG recordings via a sub millisecond local loopback UDP connection and a DAQ respectively. iEEG recordings from up to 256 electrodes (visualized in MRI) are digitized at 1 KHz and combined with digital triggers using Ripple Neuro’s Grapevine Neural Interface Processor (NIP) are transmitted and stored on a computer.

A MATLAB^TM^ script, running on the same laptop as the SMI iView ETG Server software, broadcasts numbered triggers every 10 seconds, injecting them simultaneously into the neural data stream via a Measurement Computing USB-204 data acquisition (DAQ) device connected to the NIP’s digital port and into eye–tracking event stream via SMI’s iView ETG server application via a sub millisecond latency local loop back network connection using UDP packets (Fig. 2). These triggers were used to align and fuse the heterogeneously sampled data streams after the experiment, during the *Data Fusion* stage (see below for details).

#### Best Practices for Behavioral Recording

In each recording session, neural activity recording was initiated followed by simultaneous initiation of recording of eye–tracking, egocentric video, and audio recording via the SMI ETG 2 Eye Tracking Glasses using the SMI iView ETG Software Server. Once the recording of all modalities was underway, the MATLAB^TM^ script was initiated to generate and transmit triggers. At the end of each recording session, the tear down sequence followed the reverse order: 1) the MATLAB^TM^ script was terminated, marking the end of the recording, 2) the SMI iView ETG Software Server recording was halted, 3) the neural data recording stream was stopped on the NIP. Excess data from prior to the first numbered trigger and after the last numbered trigger were discarded for all modalities.

Shift in the placement of the eye–tracking glasses is possible if the participant inadvertently touches or moves them during a recording session. Such disruption can introduce systematic error(s) in eye gaze data captured after the disruption(s), although errors can be mitigated with gaze correction (see *Data Preprocessing* for details). The potential for such an event increases with the duration of a recording session. To minimize the risk of such error(s), we first instruct participants to avoid touching or nudging the glasses during a recording session to avoid disrupting the eye–tracking calibration completed at the beginning of the recording session. Second, we strive to reduce such errors by limiting an individual recording session to one hour and including a short break for participants. During this interlude, the recording is terminated, and participants are offered the opportunity to remove the eye tracking glasses before initiation of the next session. The interlude serves two purposes: 1. it gives the participant a break from wearing the eye–tracking glasses, helping to alleviate fatigue and discomfort; 2. initiating a new recording allows the research team to re-secure and re-calibrate the eye–tracking glasses, renewing the accurate mapping of gaze to the egocentric video. Although we prefer ≈1 hour recordings as a best practice, maintaining this practice depends upon participants’ preference and the number visitors. In some cases, recording sessions may be longer.

### Ergonomic Modifications to Eye Tracking Glasses

Standard clinical care following iEEG implantation involves the application of a bulky gauze head dressing. This bandaging is applied around the head to protect the operative sites where the iEEG electrodes are secured with bolts. The dressing also includes a chin wrap to provide further support in preventing dislodgement of the iEEG electrodes by securing the connector wires that carry electrical activity to clinical and/or research recording systems like the Ripple Neuro Grapevine NIP. In our studies, the bandaging typically covered the participants’ ears, rendering the temples on the eye-tracking glasses unusable. To overcome this challenge, we modified the structure of the eye-tracking glasses, removing the temples and substituting them with an adjustable elastic band. We attached the elastic band to the frame of the eye-tracking glasses using Velcro patches sown at each end. The modification permitted secure placement of the glasses on the face of a participant, with the elastic band carefully stretched over the head dressing to avoid disturbing the operative sites (Fig 1.c). To reduce any pressure the eye-tracking glasses placed on the participants’ faces as a result of the elastic band alteration, we further modified the glasses by adding strips of adhesive backed craft foam to the nose bridge and upper rims of the frame. These ergonomic solutions enabled correct, robust, and comfortable placement of eye-tracking glasses for each participant with flexibility to adjust to individual bandaging and electrode placement configurations. As an added measure to minimize the possibility of movement for eye-tracking glasses during recording sessions, the USB cable connecting the eye-tracking glasses to the laptop was secured to the participants’ hospital gowns near the shoulder with a large safety pin to prevent the weight of the remaining length of cable from pulling on and displacing the glasses during a recording session. Sufficient slack was left in the cable segment between the glasses and the fixation point on the participants’ gowns to allow for free head movement while preventing the secured cable segment from pulling on and potentially displacing the eye tracking glasses.

### Data Preprocessing

The behavioral (eye-tracking, video, audio) and physiological (neural) data streams captured during a real-world vision recording were preprocessed as follows before *Data Fusion* was initiated.

#### Eye-Tracking

The eye-tracking data stream is composed of time series data sampled at 60 Hz, where each sample (referred to as an eye-tracking trace) contains a recording timestamp, an eye gaze location (X,Y coordinates in the space of egocentric video) and is labeled by the SMI iView ETG platform as belonging to a fixation, a saccade or a blink. Consecutive eye-tracking traces with the same label (fixation, saccade, or blink) are interpreted as belonging to a single eye-tracking ‘event’ of that type, whose duration is the difference in recording timestamps of the last and first eye-tracking traces in the block of consecutive traces with the same label (fixation, saccade or blink).

As an example, a set of 60 eye-tracking traces (amounting to 1 second of recorded activity), where the first 30 are labeled as fixation, the next 12 labeled as saccade, followed by the final 18 labeled as fixation, would be interpreted as a fixation event ≈500 ms long (30 samples at 60 Hz), followed by a saccade event ≈200 ms long (12 samples at 60 Hz) followed by a fixation event ≈300 ms (18 samples at 60 Hz).

We developed custom Python scripts that parse eye-tracking traces and construct logs of eye-tracking events for each recording session. In addition to the duration of each eye-tracking event, the median gaze location (median is used for robustness to outliers) was logged for each fixation event and the start/end gaze locations were captured for each saccade event. Blink traces are denoted by a loss of eye-tracking (i.e. absence of gaze location) and as a result only the duration of blink events was tracked in the consolidated eye-tracking event logs.

Preprocessing of eye-tracking data also incorporates the detection and correction of systematic errors in gaze angle estimation that can be induced by the movement of eye-tracking glasses during recording sessions (e.g., if a participant inadvertently touches and moves the glasses due to fatigue), which disrupts the calibration of eye–tracking glasses (see *Data Acquisition* for details). Such issues were detected by manually viewing all experimental recordings using SMI’s BeGaze application, which renders eye-gaze, audio and egocentric video together. The disruption of calibration for eye gaze tracking is visually detectable when viewing egocentric video overlaid with eye-tracking and audio because visual behavior is altered such that the gaze data fails to make sense consistently after loss of eye-gaze calibration (e.g., the subject is scrolling through a phone or reading a book or watching tv or talking to someone, but the gaze location is visibly shifted away from the obvious target). These issues were corrected using the SMI BeGaze application, which allows researchers to apply a manual correction (i.e., an offset) to eye gaze at any time point in a recording, which applies to all eye gaze data following the corrected time point. The corrections were verified by reviewing the video that followed the correction, to ensure that corrected eye gaze data made sense consistently. Corrections to eye-tracking data preceded preprocessing in such cases.

#### Video

Recordings of egocentric (head-centered) videos offer a broad range of visual stimuli, including objects, people and faces. Since the video recordings come from a camera mounted on the same glasses as the eye tracker they provide an egocentric view, i.e. the recorded videos capture the scene corresponding to where the participant is facing and the perspective moves as the participant’s head moves. As a broad research goal, we wanted to know what objects were present in the recorded scenes. Our primary object’s of interest were visitors’ faces and bodies, given the objective of examining social interactions. We processed videos to identify the location of faces and body parts of people in the video recordings. As a secondary objective, we were also interested in identifying other non-face and non-body objects. Finally, for all face locations, we extracted several higher-level measures about human visual behaviors, including head pose (including orientation and position of the head), eye gaze (e.g., toward vs away from the observer) and facial expressions.

To automatically identify faces, people, and other objects, we used a computer vision algorithm - YOLO v3 (Redmon et al., 2016) for object detection on each video frame. The algorithm identified bounding boxes and labels for each object present in a video frame, including faces and people. A total of 1,449,098 video frames were processed this way. While there has been great progress in computer vision for automated object detection in the last decade, it is not perfect. For example, algorithms such as YOLO v3 are trained on image data sets which contain a predetermined list of object categories, that may not include many objects that are present in a clinical setting. In addition, objects belonging to the predetermined list of object categories may also be mis-detected (false positives or false negatives). Since the annotations were supposed to serve as ground truth for analysis of neural data, their accuracy was essential and we implemented a second stage of annotation based on human judgement, to confirm the quality of automated object detection and correct mis-detection. To avoid the time intensive prospect of manually annotating all video frames in the second stage, we annotated the first video frame corresponding to each fixation, because fixations are typically brief (a few hundred milliseconds), defined by the lack of significant eye movement, and thus it is reasonable to assume that participants look at the same location/object in a relatively unchanging scene during a fixation. We identified the video frame corresponding to the beginning of each fixation using video timestamps present in eye-tracking traces. Human annotators provided coordinates of bounding boxes for each face, or person present in video frames for a total of 125,996 frames as part of the second stage of annotation.

Finally, we used the OpenFace software (Baltrusaitis et al., 2018), a facial behavior analysis toolkit using computer vision technology, to extract additional high-level information for face regions. For each face region, OpenFace provides information about (1) the position of 64 facial landmarks including eye, nose and mouth positions, (2) head orientation and position, (3) eye gaze direction and (4) facial expression information encoded following the Facial Action Coding System standard (Friesen and Ekman, 1978).

#### Audio

Audio recordings from a microphone embedded in the eye–tracking glasses capture sound from the participant’s perspective. The clarity of recorded audio is influenced by the loudness of sounds and the distance of the source from the participant. Since our objective involves examining social interactions, speech detection and speaker identification are ”events” of interest.

To detect time segments with speech in the audio recording and to diarize the audio (i.e. to determine who spoke when) we use state of the art deep learning speech detection (Lavechin et al., 2020) and speaker identification (Yin et al., 2018) pipelines available as part of an open source toolbox (Bredin et al., 2020). Even these state-of-the-art models have unacceptably high error rates (particularly for diarization) for them to provide useful annotations as labels in analysis of behavior-physiology relationships. In order to overcome this hurdle, we configured these models to be highly sensitive (leading to higher false positives, but very few false negatives) and then manually reviewed model predicted time segments for speech and speaker identification, to identify and correct false positives. Outside of parameters that control the sensitivity of the deep learning models, the efficacy of speech detection and diarization is influenced by the loudness of the speakers themselves, as well as their distance from the participant (i.e, the microphone). This means that the participant’s speech is the most reliably detected, while the quality of speech detection (and therefore speaker identification) for other speakers may vary. As a result, we chose to collapse audio diarization into two categories during manual review, the participant and speakers other than the participant. Segments with concurrent speech from the participant and other speakers were labeled as participant speech.

#### Intracranial Recordings

Response potentials and broadband high frequency activity (BHA) were extracted from the raw iEEG recordings for statistical analysis using MATLAB^TM^. Response potentials were extracted using a fourth order Butterworth bandpass ([0.2 Hz, 115 Hz]) filter to remove slow linear drift and high-frequency noise, followed by line noise removal using a fourth order Butterworth bandstop ([55 Hz, 65 Hz]) filter.

BHA extraction involved two steps. First, the raw signal was filtered using a fourth order Butterworth bandpass ([1 Hz, 200 Hz]) filter followed by line noise removal using notch filters at 60, 120 and 180 Hz to obtain local field potentials. Next, Power spectrum density (PSD) between 70–150 Hz was calculated for the local field potentials with a bin size of 2 Hz and a time-step size of 10 ms using Hann tapers. For each electrode, the average PSD across the entire recording was used to estimate a baseline mean and variance of the PSD for each frequency bin. The PSD was then z-scored using these baseline measurements for each frequency bin at each electrode. Finally, BHA is estimated by averaging the z-scored PSD across all frequency bins (excluding the line noise frequency bin at 120 Hz).

iEEG recordings were subjected to several criteria for inclusion in the study. Any recordings with ictal (seizure) events were not included in the study. Artifact rejection heuristics were implemented to avoid potential distortion of statistical analyses due to active interictal (between seizure) or outliers. Specifically, we evaluated the filtered iEEG data against three criteria that are applied to each sample i.e., each time point in iEEG recordings, which corresponds to 1ms of neural activity. These criteria were applied to the filtered iEEG signal for each electrode, as well as the averaged (across all electrodes) iEEG signal. The first criterion labels a sample as ‘bad’ if it exceeds 350 *µ*V in amplitude. The second criterion labels a sample as bad if the maximum amplitude exceeds 5 standard deviations above/below the mean. The third criterion labels a sample as bad if consecutive samples (1 ms apart at a 1000 Hz sampling rate) change by 25 *µ*V or more. For the averaged iEEG signal, any sample satisfying any of these three rejection criteria is labeled as bad. Further, if more than 10 electrode contacts (out of a typical 128) satisfy the bad sample criterion for a particular sample, it is labeled as a bad sample. Less than 10% of the samples in experimental recordings were labeled as bad samples. All data types were dropped from analysis for fixations that contained bad samples.

### Data Fusion

Precise fusion of heterogeneous behavioral (eye-tracking, egocentric video and audio) and physiological (neural) data streams is essential for constructing a multimodal data set to answer our questions about the neural correlates of real-world vision. In our approach, eye-tracking provides the reference modality against which video/audio, psychophysiological, and neurophysiological (neural activity) data streams are aligned in time (Fig 3). Each eye-tracking event is mapped to a corresponding egocentric video frame. For fixation events, we combine eye gaze location with bounding box locations/sizes from annotations for the egocentric video frame to determine what object (face or non-face) the participant is fixating upon. Each eye-tracking event is mapped to an auditory time segment and labeled as belonging to a speech or silence segment, with additional labeling for speaker identity in the case of a speech segment. Finally, neural recordings are also aligned in time to eye-tracking events based on the temporal offset of eye-tracking events and neural data, from trigger events which are injected in both data streams at 10 second intervals during recording sessions.

**Figure 3:**
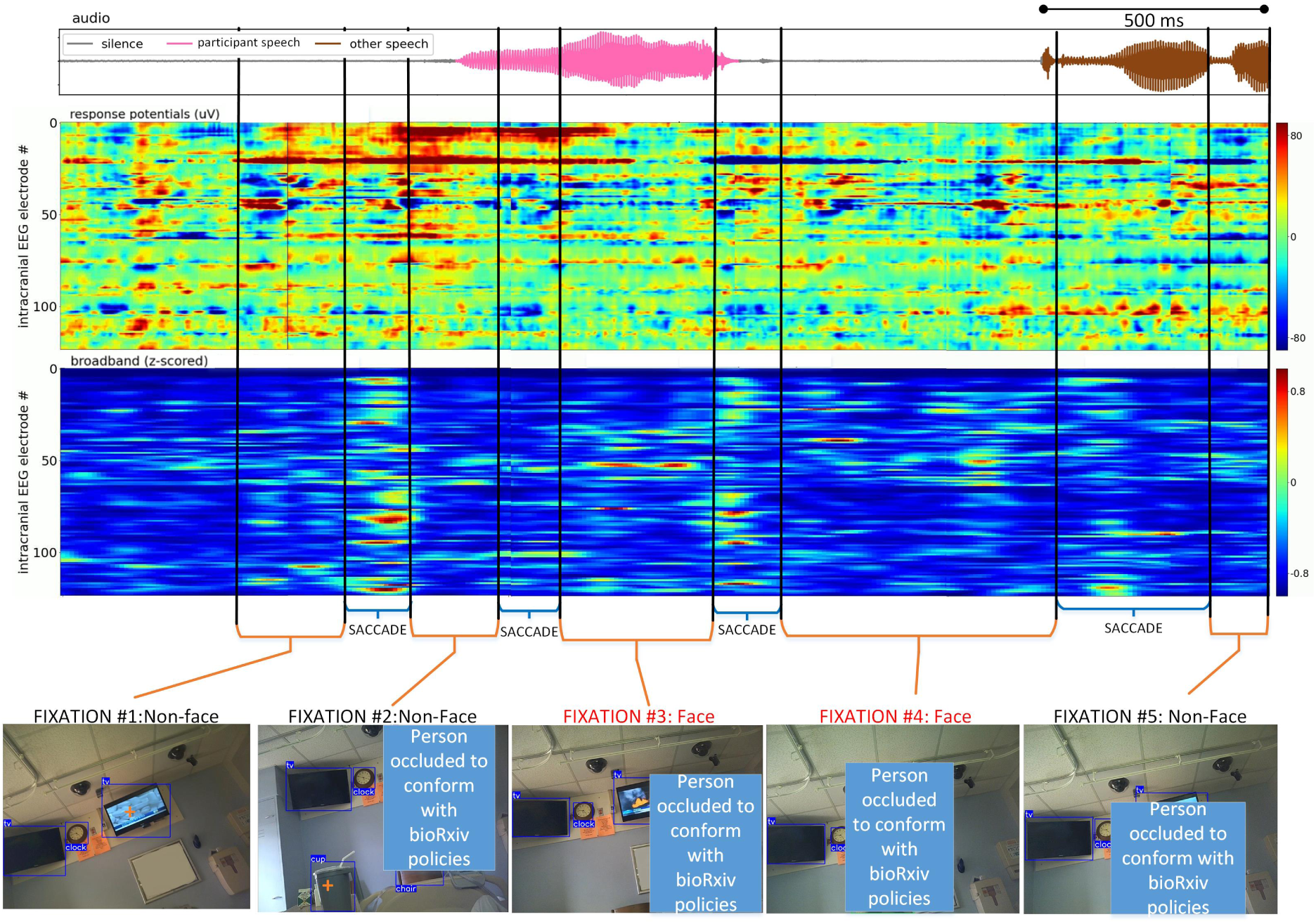
Fused multimodal data set from a real world vision recording: The audio waveform is shown on top, with gray, pink and brown segments denoting silence, participant speech and speech from other speakers. Response potentials and broadband high frequency activity heat maps from a 124 iEEG electrode montage are shown below the annotated audio. Vertical black lines demarcate fixations and saccades, which are marked underneath the audio and neural time series with orange and blue braces respectively. The bottom row shows video frames corresponding to each fixation event, with an orange ‘+’ denoting eye gaze location and bounding boxes identifying the location of different objects, including persons and faces.

The quality of multimodal data sets assembled by the data fusion process described above is reliant on the quality of the heterogeneously sampled behavioral, psychophysiological, and physiological data streams fed into the data fusion process. Acquisitional variability, if present and left undetected, can severely degrade the quality of fused data sets by introducing alignment issues, and dropped video frames and/or recording offsets are common. Our methodology includes cross-verification procedures that guard against such issues with careful examination of the raw data streams for each modality. These procedures assume that the raw data captured for any modality contains accurate and sufficient timing information to diagnose and correct such issues. As long as hardware/software systems in use meet this basic assumption about raw data capture, the cross-verification approach we describe should scale. Below, we detail two specific issues that arose in our recordings using SMI ETG 2 Eye Tracking Glasses and illustrate how we addressed them to ensure data quality in the fused data set.

#### Sampling rate variability

Variability in sampling rates is observed in engineered systems and can arise due to a variety of reasons ranging from temperature dependent variation in the frequencies of crystal oscillators that drive digital clock signals to propagation delays in circuit boards and circuitry running at different clock rates. If a fixed sampling rate is assumed, then these variations can accumulate as sampling drift over time and potentially lead to significant timing offsets over long periods of operation. These phenomena are addressed in engineered systems in various ways including using clocks far faster than the sampling rates desired and frequent resetting/calibration to minimize drift accumulation.

Here, we describe our approach to detect and remove such issues from the final multimodal data set that results from our data fusion procedures. We evaluated variability in the sampling rate of eye–tracking traces based on their timestamps. Since audio, video and neural data are anchored to eye–tracking events, minor sampling variability for eye–tracking does not introduce any error as long as other data streams can be aligned to eye–tracking correctly. We evaluated the timing and mapping of all other modalities (audio, egocentric video and neural data) against eye–tracking. Specifically, we found the need to address sampling rate variability that arose in the egocentric video stream, so it could be reliably mapped to eye–tracking data.

The inter-frame interval for the video stream can vary systematically by small amounts from the rated 41.66 ms (24 fps) for a recording session. These deviations can be a source of error in the mapping of eye–tracking traces to video frames unless they are addressed during data fusion. A critical marker of this problem is an inconsistency between the number of frames present in the video and the number of video frames estimated from eye–tracking traces using Eq 1. It is important to note that this variability is not always accounted for in manufacturer software or documentation. The solution to this issue is relatively simple because the eye–tracking traces include a ‘Video Time’ column which has millisecond resolution. Instead of assuming a fixed frame rate as Eq 1 does

We estimated video frame numbers corresponding to each eye-tracking trace using the ‘Video Time’ in them as follows

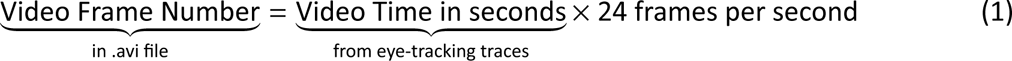

#### Addressing data gaps or corruption from behavioral modalities

Loss or corruption of data during media recordings on embedded devices is demonstrable, and is a potential source of error for a fused multimodal data set that relies on precise alignment of multiple heterogeneously sampled modalities. As a result, our data fusion process pays close attention to identifying and characterizing such issues and addressing them to ensure data integrity. Here, we qualitatively describe different classes of issues observed in our data and how we address them to ensure data quality.

We observed missing frames in the egocentric video stream. Specifically, after correcting for sampling rate variability, we observed residual discrepancies between the number of frames that were expected per the video timestamps in the eye-tracking logs and the number of actually frames present in the video files from recordings. By evaluating timestamps for each frame in the ‘.avi’ files using OpenCV (Bradski, 2000), we found that the lost frames were at the beginning of the video stream (i.e., the first *K* frames of an *N* frame video are missing) frames. We confirm this diagnosis with an additional form of verification, which used low level audio and video processing tools to manually blend audio and video streams with and without a correction for missing frames and visually verifying the absence of lip-audio synchronization issues in the resulting video. Finally, we obtained an additional point of manual verification by visualizing the ostensibly lost frames (decoders discard frames they deem corrupt when parsing a file, but they are present in the files) from the video file on a frame by frame basis, confirming that they are corrupted/garbled. The specific pattern of error (first *K* frames missing) observed with our experimental equipment (SMI ETG 2 Eye Tracking Glasses) may not replicate with other hardware, though given engineering constraints, other errors may arise instead. As an example, other eye-tracking glasses may have frame loss/corruption intermittently during a recording instead of at the beginning. However, our observations suggest that such issues may exist with other eye-tracking devices and data fusion pipelines should incorporate verification stages that can identify and correct such issues, with a preference for multiple modes of verification that are consistent with each other.

Blinks are a natural part of visual behavior and the eye-tracking records denote them as such. Since eye–tracking is lost during blinks, there is usually no information about gaze, pupil dilation etc. available for blink events. We see blinks interspersed among fixations and saccades, and they are typically a few hundred milliseconds long. However, we observed longer periods lasting several seconds in multiple recordings. To understand this phenomenon better, we viewed the videos for periods where this happened, with gaze information overlaid using SMI’s BeGaze software. We found these anomalous blinks to be masking a real phenomenon, where the participant may be looking out the corner of their eye, which takes their eye-gaze outside of the field of vision of the egocentric camera or upon occasion, potentially taking their pupils outside of the field of vision of the eye–tracking camera. Since the system cannot accurately capture visual behavior as it relates to the video in these conditions, it labels those periods as blinks. These scenarios are easy to spot during manual inspection because the eye-gaze location before and after the blink tends to be near the periphery of the video frame. These conditions are not a significant challenge for data quality, because they can be easily dropped from analysis. However, awareness of their existence is meaningful for data fusion pipelines.

## Results

We collected iEEG recordings from patients in the Epilepsy Monitoring Unit (EMU) who wore SMI ETG 2 Eye Tracking Glasses as they went about their day interacting with friends and family visiting them as well as members of the clinical team. We used computer vision models to identify objects, faces and persons (bodies) in videos of the visual scenes in front of the participants during these sessions. Similarly, we used speech processing models to identify speech intervals and diarize the audio recorded from the internal microphone in the SMI ETG 2 Eye Tracking Glasses. All annotations from computer vision and speech processing models were validated and corrected, if necessary, by human annotators to ensure data quality. Here, we show that fused multimodal datasets (see Fig 4 for a snapshot; see *Supplemental Video* for a dynamic version) which include annotated audio, eye-tracking, annotated video, and iEEG, can be developed using this process. Such datasets can help advance visual neuroscience research beyond traditional experimental paradigms and explore the neural correlates of real-world social vision.

**Figure 4:**
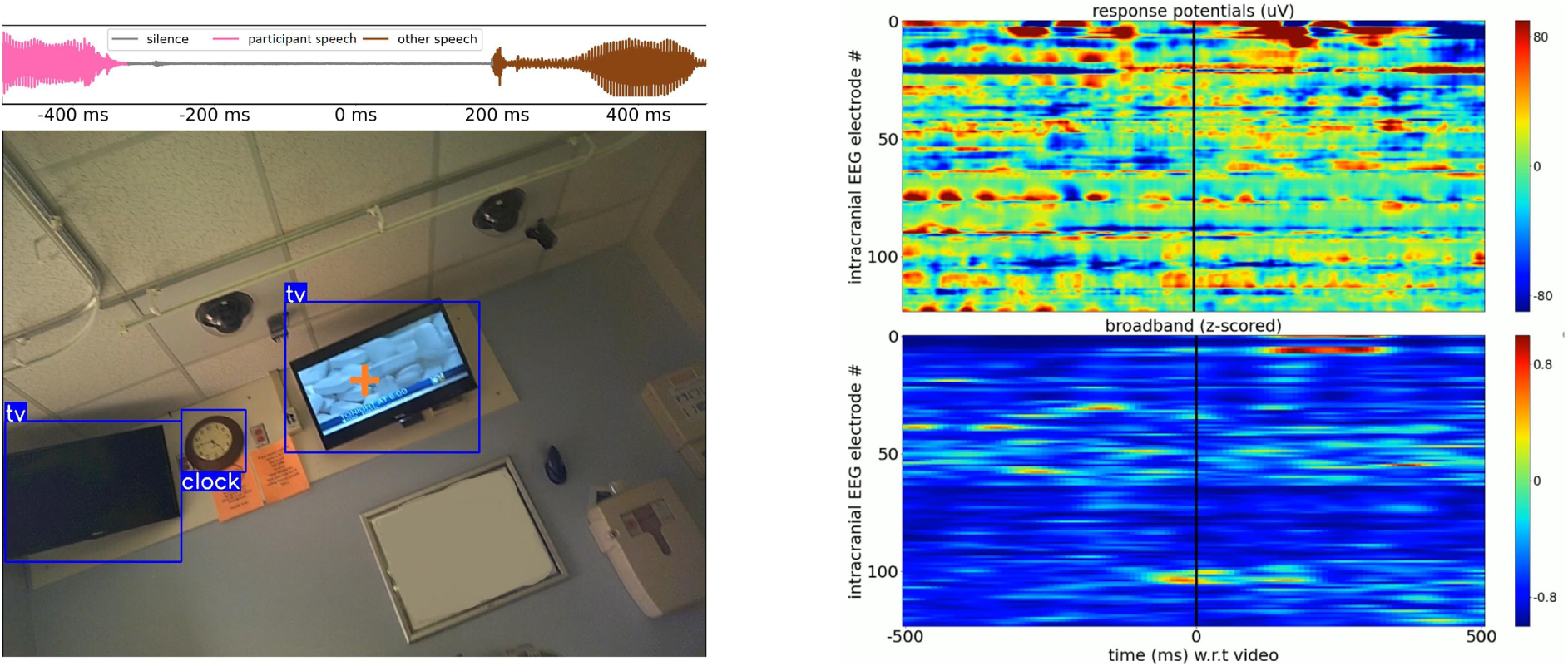
A snapshot of fused multimodal (audio, egocentric video, eye-tracking and iEEG): On the left, an annotated audio snippet (top) and video frame (bottom) visualizes the world through the participant’s eyes and ears as they interact with friends and family visiting them during a recording session. Speech/silence and speaker diarization labels color the audio signal on top. The annotated video frame below depicts the participant’s eye gaze location with orange ‘+’ marker with colored bounding boxes identifying the location and sizes of different objects detected by computer vision models and verified by human annotators. The panel on the right visualizes 1 second of neural activity across 124 iEEG electrodes, corresponding to the video frame/audio on the left, with response potentials on top and broadband activity at the bottom (see *Supplemental Video* for a dynamic version of this figure).

### Behavioral Data

We collected data from 6 participants across 11 different free viewing recording sessions which ranged from 41 - 143 minutes long and added up to a total of 16 hours and 48 minutes. Social contexts differed across recording sessions and sometimes within a recording session, in terms of the number of individuals present, the number of interactions they had with the participant and the nature of those interactions.

#### Visual Behavior

SMI Eye Tracking glasses captured visual behavior, categorizing each moment’s sample as belonging to a saccade, fixation, or blink. Visual behavior varied depending upon the social context during recording sessions. Saccades usually accounted for 10 - 15% of the recording duration (Fig 5.a), even though they account for nearly half the events (after accounting for blinks and occasional loss of eye-tracking) (Fig 5.b) as a result of the saccade–fixation–saccade structure of the active sensing cycle, a contrast highlighted by the skew in the distribution of saccade durations and fixation durations (Fig 5.c). Saccades and fixations are not perfectly balanced due to the loss of eye-tracking from blinks and other reasons (e.g., noisy conditions, participants closing their eyes for brief periods or looking out of the corner of their eye during the recording sessions).

**Figure 5:**
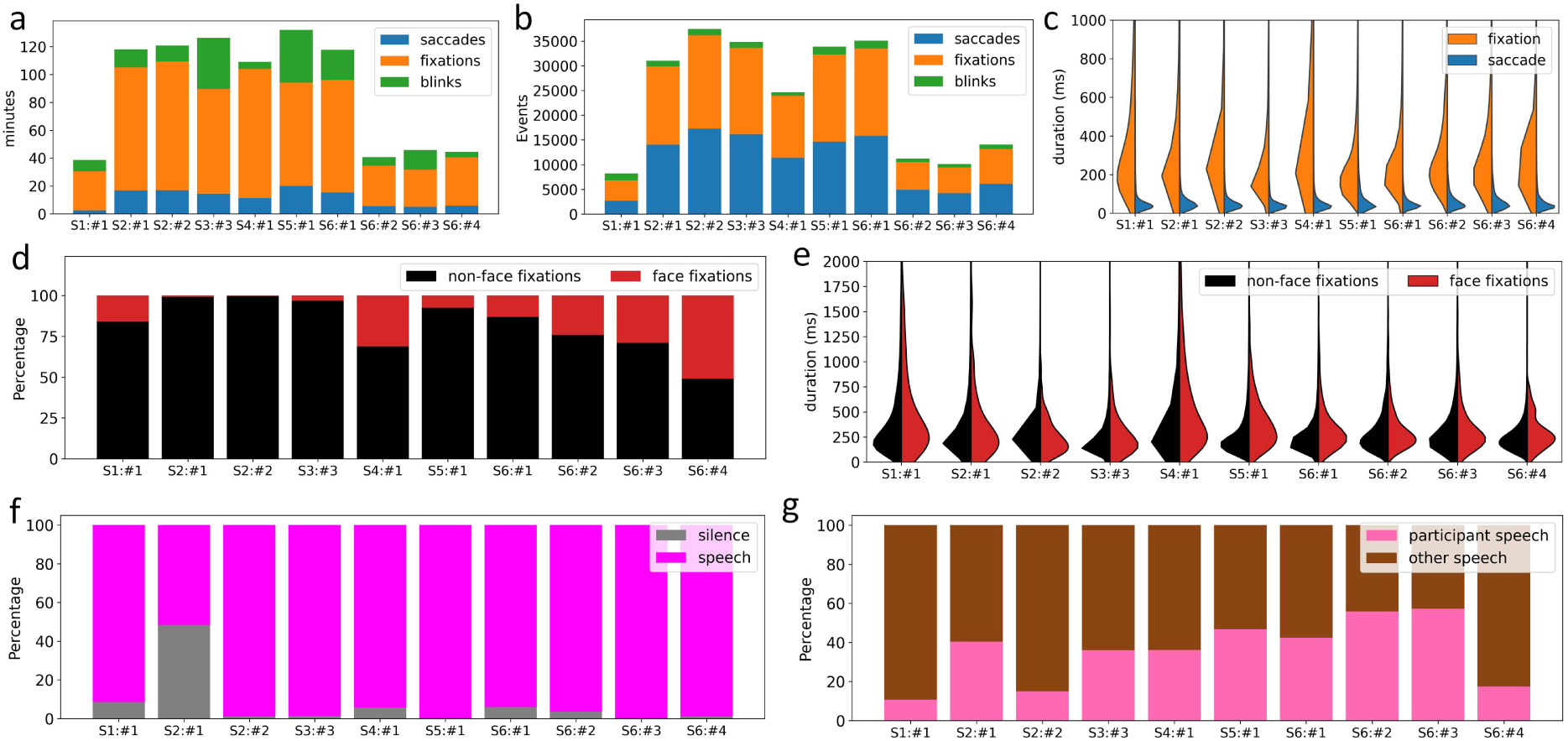
Summary of dataset spanning visual behavior and the auditory environment: **a)** The duration of each recording session (with multiple sessions for each participant) broken down by time spent in different visual behaviors (saccades, fixations and blinks). **b)** Similar to a), but counting distinct events for each visual behavior instead of time. **c)** Saccade and fixation duration distributions for each recording session. **d)** The fraction of time fixations were on faces and non-face objects for each recording session. **e)** Fixation duration distributions for face and non-face targets for each recording session. **f)** The fraction of each recording session broken down by time spent in silence and speech. **g)** The fraction of each recording session broken down by speech from the participant and other speakers.

We identify fixation targets by combining gaze location from eye-tracking with bounding boxes from the video frame corresponding to each fixation. We categorize fixations as face and non-face fixations, reflecting our focus on the social aspects of real-world vision. The social context during a recording session has a natural influence on the distribution of fixation targets. We found that participants fixated on faces less than 30 - 40% of the total time spent fixating during a recording session (Fig 5.d), even in the most social situations (e.g., EMU room full of multiple family and friends, with active conversations). The distribution of fixation durations for the two fixation categories showed that face fixations tend to be a little bit longer (Fig 5.e), indicating that even during the most social situations with familiar people, we look at the faces of people around us infrequently but when we do look at them we tend to hold them in our gaze a little longer.

#### Auditory Context

The SMI ETG 2 Eye Tracking glasses also recorded audio using an in-built microphone. We used deep learning models (Bredin et al., 2020) to do auditory scene analysis, augmenting it with manual annotation to ensure high reliability. Once again, depending upon the social context during each recording session, we observed varying levels of verbal discourse (Fig 5.f). We observed that speech could be detected from both the participant and others in the room, but the participant was reliably comprehensible due to their proximity to the microphone, whereas the volume and comprehensibility of the voices of other speakers would vary based on how close they were to the participant, making source identification more challenging even for manual annotation. To avoid potential confusion during manual annotation, we restricted speech diarization during supplemental manual annotation/verification to classifying speech as originating from the participant or other persons in the room. We found that the participant’s own verbal behavior varied across recording sessions, with comparable speech in the room, across recording sessions, even for the same participant (Fig 5.g).

#### Behavioral Annotation: Reliability and its Cost

##### Egocentric Video

Automated software driven annotation of video frames is straightforward and fast, but accompanied by a trade-off between speed and accuracy. The speed of automated annotation depends upon the algorithms used for object detection. YOLO v3 (Redmon et al., 2016) is a popular algorithm for object annotation (detection), performing at a rate of 45 fps on a NVIDIA K40 Graphics Processing Unit (GPU), or 5 fps on a standard CPU. This means annotating an hour of video takes 32 minutes with a GPU, or close to 5 hours with a CPU.

The accuracy of annotation algorithms is not high. We measured the quality of the automated annotations by comparing the automated annotations from software with human annotations for all sessions. We found that software-driven annotation only achieved an average of 69.5% Intersection over Union score (a measurement for evaluating object detection algorithms, higher the better, with a threshold of 100%). This means that the overlap ratio between the software’s bounding boxes and the human annotators’ bounding boxes was only 69.5%, suggesting the accuracy of automated software-driven annotation may be limited.

Although human annotators produce higher-quality annotations, the process is time and labor-intensive. Human annotators annotated 125,996 frames out of 16 hours and 48 minutes of videos. The total time spent on annotating 125,996 frames was 104 hours, with an average of 6.19 hours for an experienced human annotator to annotate an hour of video.

For quality control, 3% of frames from two sessions (sessions from S5:#1 and S6:#4) were randomly sampled and verified by a second annotator. The overlap ratio between the first annotator’s bounding boxes and the second annotators’ bounding boxes was 97.3% and 97.4% respectively for the two sampled sessions. This process underscored the significantly higher quality of human annotation over the automated software-driven annotation.

##### Speech Detection and Diarization

Automated speech detection and speaker identification were computationally efficient, with an hour’s audio being processed within 1 - 2 minutes. Manual verification and correction of mis-detection was done with manual annotator’s listening to the audio, and correcting false negatives/missed speech and false positives/speech labeled as silent and required an hour of manual effort for each hour of audio recording. Comparisons of automated speech detection with manually verified/corrected speech intervals for the first 10 minutes of each recording session revealed mis-detection (speech classified as silence or vice versa) for ≈ 4% of the annotated audio. Manual annotation for speaker identification involved collapsing the automated speaker diarization labels into two categories, ‘participant speech’ and ‘other speech’. Speech segments where the participant and other individuals were speaking concurrently were labeled as ‘participant speech’. Manual annotators listened to the full length of the recording assigning new labels to each speech segment manually, which took 75 minutes for each hour of speech.

#### Data Fusion Issues: Detected and Corrected

Next, we show some results which motivate careful evaluation of the raw data for each modality before data fusion of heterogeneously sampled data streams from an experimental recording is attempted. Specifically, we describe and quantify alignment issues between eye-tracking and video data collected using SMI ETG 2 Eye Tracking glasses, that were identified and corrected (see *Methods for details)* during data fusion. We found two issues in the video stream, which would lead to misalignment between eye-tracking traces and the video frame they correspond to.

The first issue was related to corrupted and unrecoverable egocentric video frames at the beginning of each recording (see *Methods* for details). The duration of egocentric video lost as a result of this issue varied by recording, and ranged from the first 0 ms - 625 ms (Fig 6.a). In a video with the first *N* frames corrupted, this issue would lead to incorrect mapping of eye-tracking traces to a video frame *N+1* frames later than the egocentric video frame they corresponded to, which could lead to errors in annotation of fixations (e.g., as face or non-face fixation) across the entire video. After correction, the only impact of this issue is that eye-tracking traces/neural data for the first few frames that are corrupted and discarded cannot be used for analysis, which is a very minor loss.

**Figure 6:**
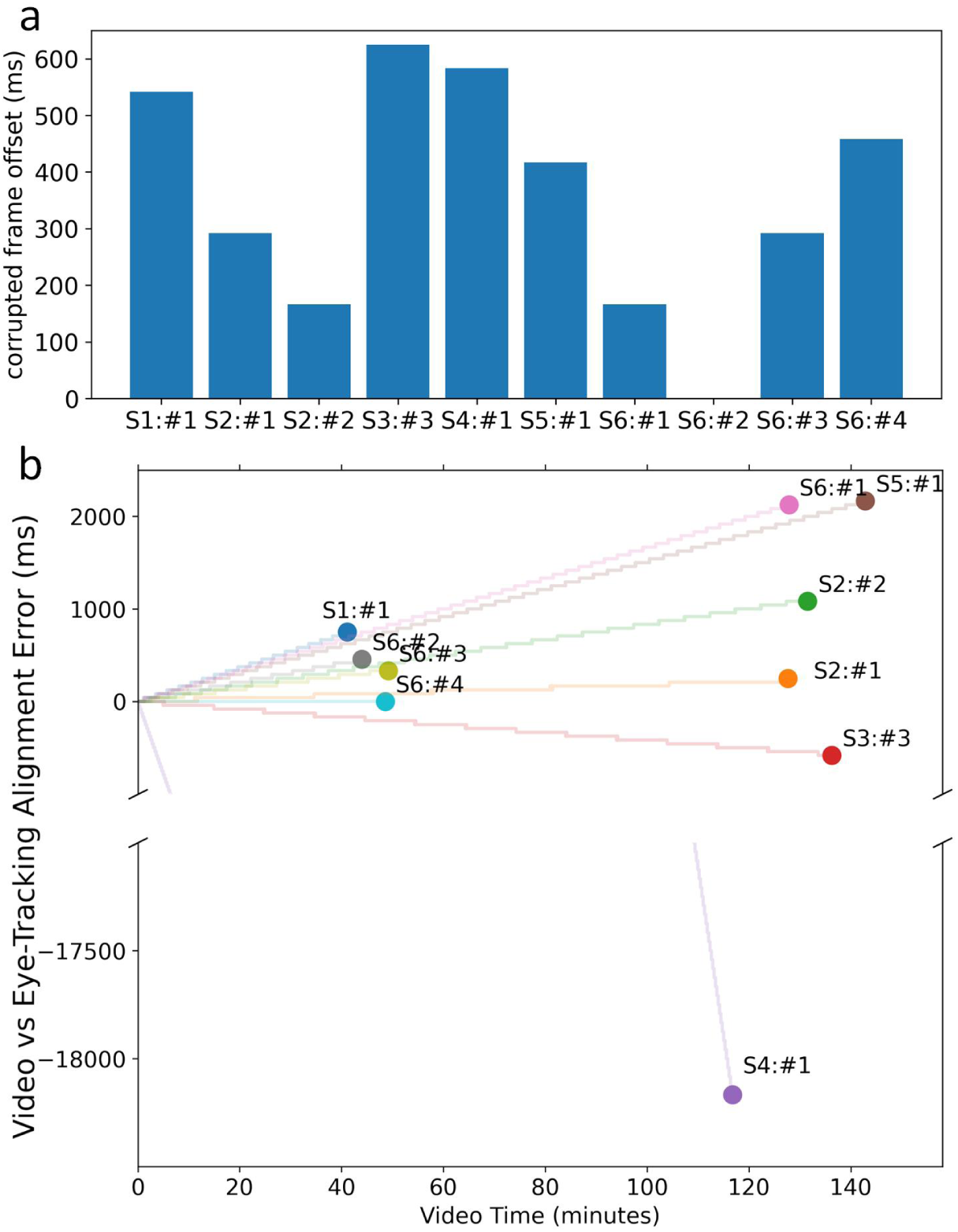
The potential effects of video frame corruption and video frame rate variability on the accuracy of data fusion: Visualization of timing error for each recording session introduced in the alignment of eye–tracking events and corresponding egocentric video frames in the case **a)** Corrupted frames at the beginning of each video file are not detected and corrected in the eye–tracking to video frame alignment procedure. This is a fixed error that affects all eye-tracking events in a session. **b)** The procedure to map frames to eye–tracking traces does not address small variations in frame rate (i.e., Eq. 1 instead of Algorithm 1). This is a time varying error which accumulates over the duration of a recording (scatter points indicate the final accumulated error at the end of each recording session) and its rate of accumulation (slope of shaded lines in the background) depends upon the magnitude of the deviation in video frame rate is from 24 fps. We observe deviations as small as 1 frame (41.67 ms) over a 49 minute recording for S6:#4 and as large as 432 frames (18 seconds) over a 2 hour recording for S4:#1.

The second issue was related to variability in the average frame rate for egocentric video recorded from each session. We observed that for different sessions, the average frame rate of the recorded video was slightly above or below 24 frames per second. Eye-tracking traces are mapped to video frames using a ‘Video Time’ variable embedded in them. Estimating the video frame number corresponding to an eye-tracking trace using Eq. 1 which assumed a frame rate of 24 fps that was slightly higher or lower than the real frame rate of the video. The discrepancy led to an error between the estimated frame and the real frame corresponding to eye-tracking traces, which accumulated as the video recording progressed (Fig 6.b) and became visible with the eye-tracking traces mapping to far fewer/greater frames than were present in the video at the end of the recording. This problem was avoided by using the procedure defined in Algorithm 1, which is robust to these small variations in frame rate (see *Methods* for details). Both these problems co-occurred and addressing them as described in the *Methods* section gave us perfect consistency between the number of frames estimated in the eye-tracking traces and the number of frames present in the egocentric video. Lastly, we also evaluated audio and neural activity for similar alignment inconsistencies with the eye–tracking logs and found no issues with alignment.

##### Algorithm 1 Sampling rate variability resistant mapping of video frames to eye-tracking traces

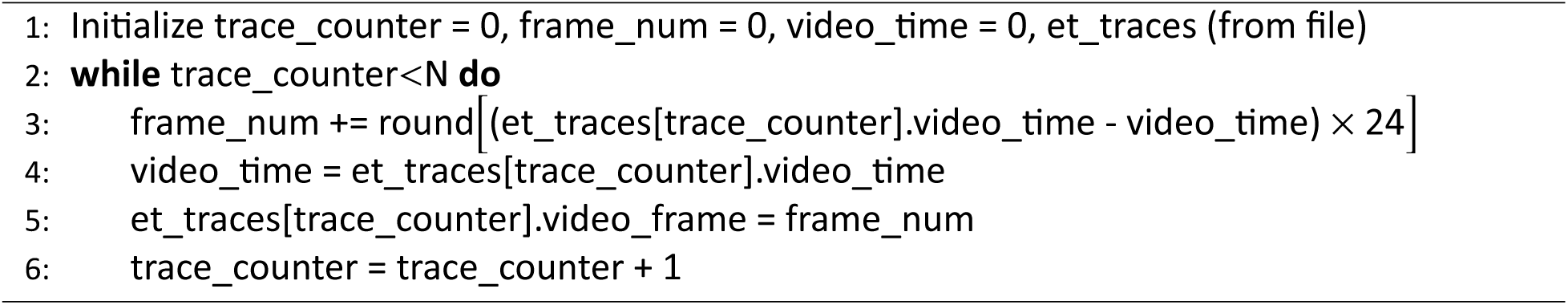

### Neural Correlates of Real-World Social Vision

The number and cortical locations of intracranial EEG electrodes from which neural data were recorded varied by participant with a total of 686 cortical locations distributed across the temporal, parietal, occipital, frontal and cingulate areas of participants (Fig 7.a, b).

**Figure 7:**
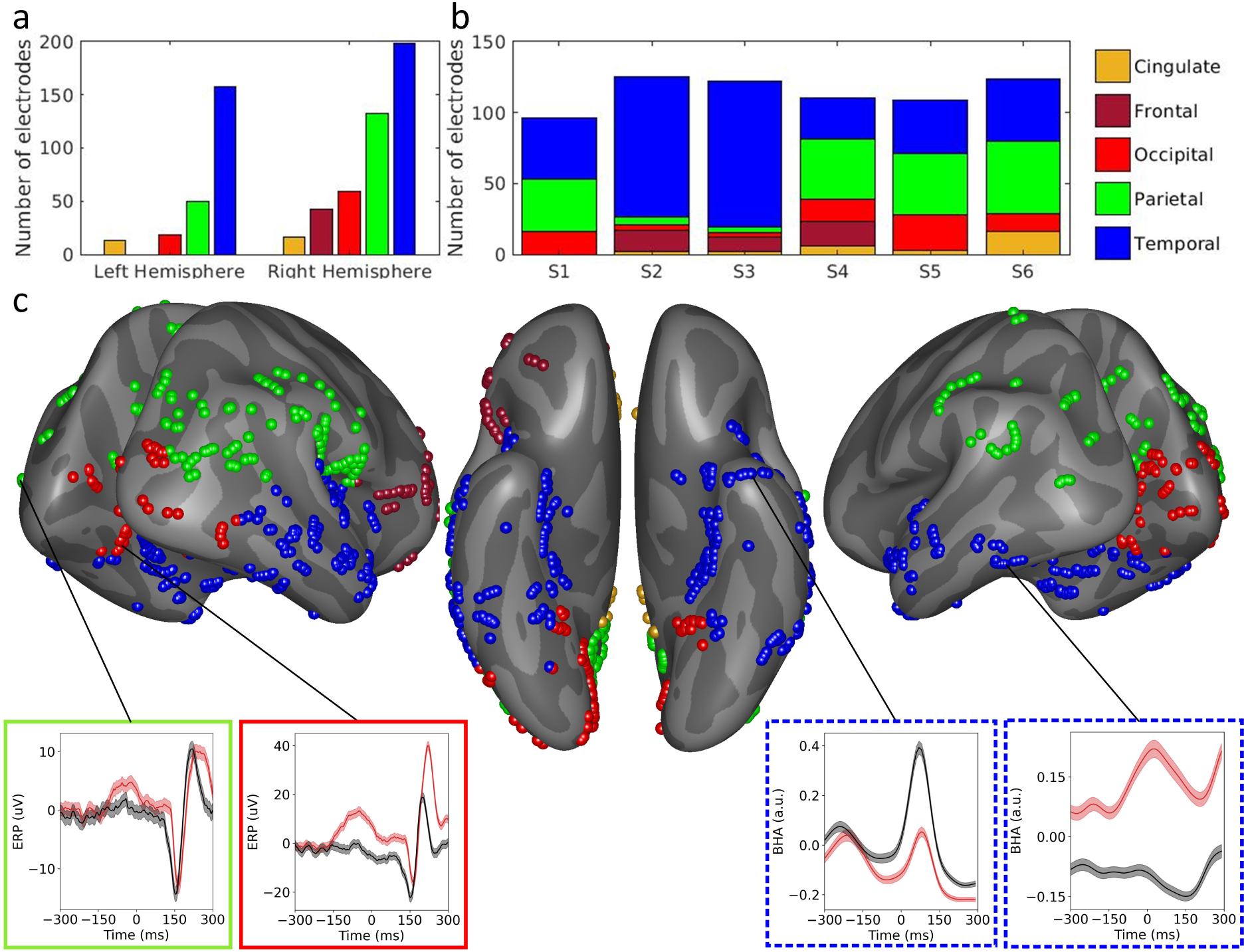
**a)** Cortical distribution of the 686 intracranial EEG electrodes from 6 participants over different lobes across the Left and Right Hemispheres per the Desikan Killiany atlas (Desikan et al., 2006). **b)** Per participant electrode distribution across different cortical regions **c)** Visualization of locations of electrodes from all participants on an inflated cortical surface with ventral, lateral (left and right), posterior and anterior views. Average fixation locked neural activity from electrodes sampled across all participants and recording sessions. The colors of the boxes correspond to the lobe of the cortical location being sampled and the outlines denote the neural signal that is visualized (solid lines denote Fixation Response Potentials (uV) and dashed lines denote Fixation Response Broadband Activity (a.u.)). The average fixation locked response to ≈1000 fixations of face (red) and non-face objects (black) each is shown for each cortical location. One notable result is that differences in the neural response between face and non-face fixations appear prior to fixation onset, suggesting predictive activity/”pre-saccadic preview”(Buonocore et al., 2020; Huber-Huber and Melcher, 2021)

Finally, we aligned neural activity recorded from intracranial EEG electrodes to the composite behavioral (eye-tracking + visual behavior + auditory context) log using digital triggers embedded in the neural and the eye-tracking data streams. This final step allows identification and extraction of neural activity corresponding to individual eye-tracking events (saccades, fixations, and blinks).

Our analysis of real-world vision is anchored to fixations, and Fig 7.c visualizes average Fixation Response Potentials (FRPs) and Fixation Related Broadband High frequency Activity (FRBHA) for face and non-face fixations from several of the 686 intracranial EEG electrodes for which real-world vision data were collected. Typical aspects of the FRP (e.g. enhanced N170 for faces, particularly in ventral temporal cortex locations) and FRBHA (Allison et al., 1999; Boring et al., 2021; Ghuman et al., 2014; Jacques et al., 2019; Li et al., 2019) are well represented for electrodes from multiple lobes suggesting the alignment of neural activity and eye–tracking events is robust and provides a key ”proof-of-principle” for this real-world paradigm, similar to that provided by recent studies in macaque monkeys engaged in free viewing of natural scenes (Barczak et al., 2019).

## Discussion

We investigated the feasibility of combining neural recordings from iEEG electrodes with eye-tracking, video and audio recordings collected using eye-tracking glasses and annotated using computer vision and speech models to generate robustly fused multi-modal data sets from unscripted recording sessions in an inpatient hospital environment. Fusion of visual behavior with neurophysiological recordings enables investigation of the neural correlates of real-world social vision and affective perception. Summary views of the data highlight the heterogeneity that emerges in uncontrolled behavior in ecologically valid settings, and underscore the need for care when trying to assess generalizability of observed effects across individuals. A natural approach to address these challenges is to define summary variables or learn then using data driven approaches like multiset canonical correlation analysis (Nielsen, 2002). The efficacy of our methodology is validated in the context of real-world social vision by fixation locked neural activity (FRPs and FRBHA) for face and non-face fixations from ventral temporal electrodes, which show category selective neural signatures that are also observed in traditional visual neuroscience experiments. Our initial findings also point to several potential opportunities for the enrichment of behavioral and physiological data collection as well questions of significant interest for clinical and translational research.

### Enriching behavioral monitoring

#### Higher fidelity capture of visual behavior

From analyzing the data sets presented here, three natural opportunities to improve the capture of visual behavior are apparent. The first entails higher fidelity data acquisition for behavioral data streams that we already capture. The eye-tracking glasses used in this study feature a single head-centered perspective (egocentric) video camera operating at 24 frames per second with a resolution of 1280 x 960 pixels capturing a 60^◦^ (horizontal) by 46^◦^ (vertical) region of the field of vision, with 2 eye-tracking cameras operating at 60 Hz. Increasing the spatial resolution of the video camera in pixels, improving the temporal resolution of both eye-tracking and video and capturing a larger fraction of the field of vision can aid in better tracking of visual behavior over a more complete portion of the field of the vision. The second opportunity requires adding a new data modality (head position) using an Inertial Measurement Unit (IMU), that can provide tracking for the physical frame of reference corresponding to each video frame. The third opportunity involves considering the addition of depth perception information for eye-gaze, which may potentially be supported by the addition of a second egocentric camera or LIDAR (Roche et al., 2021). A review of available research grade hardware (Cognolato et al., 2018) provides an account of the capabilities of several research grade devices, which can be evaluated for their suitability with respect to each of these possibilities.

#### Aural scene capture

Analysis and annotation of the auditory scene recorded using the in-built microphone embedded in the eye-tracking glasses reveals the potential advantages of capturing the aural scene as well as the limitations of having a single microphone physically attached to the patient. The potential addition of high definition microphone arrays in the room can enable a complete recording the auditory scene, including the capture and source localization of all sound, including speech. In the context of social behavior, such an enriched capture offers the opportunity to go beyond speech and speaker detection and into speech recognition, and its conversion to text (Hannun et al., 2014; Ren et al., 2019; Zhang et al., 2018) thereby allowing the use of language models that could add an additional behavior modality for semantic and sentiment analysis (Kiritchenko et al., 2014).

#### From monitoring visual behavior to visual monitoring of behavior

Heavily monitored inpatient hospital environments like an EMU are typically equipped with cameras that allow clinical care teams to monitor patient behavior. The same video streams also capture the physical behavior of other individuals (e.g., doctors, nurses, family) who are present. These video streams hold the potential to add two additional behavioral modalities to the multi-modal data set we have described. The first modality is affective behavior, for the patient and other individuals present, extracted using facial analysis tools like OpenFace (Baltrusaitis et al., 2018). The second modality is physical behavior using tools like OpenPose (Cao et al., 2019) and DeepLabCut (Insafutdinov et al., 2016; Lauer et al., 2021; Mathis et al., 2021, 2018; Nath* et al., 2019), which may enable us to explore the relationship between physiology and behavioral phenomena like interpersonal synchrony (Delaherche et al., 2012).

### Enriching physiological monitoring

As part of standard care, inpatient hospital environments feature the monitoring of a wide variety of physiological data like EKGs, EMGs, heart rate, pupillometry, blood pressure, neural recordings, pulse oximeter readings, saliva samples, urine samples as well as clinical events. A richer physiological data set than the one presented here – one that contains a greater number of the physiological modalities – can combine powerfully with behavioral markers to allow pursuit of highly relevant clinical and translational research questions.

As an example, attention and arousal are thought to be modulated by the locus coeruleus-noradrenergic (LC-NE) system. Pupil size (Alnæs et al., 2014; Murphy et al., 2014, 2011) in absence of lighting change and heart rate (Azarbarzin et al., 2014) are both considered proxies for locus coeruleus (LC) activity. A data set that fuses EKG and pupillometry with human intracranial EEG along with visual behavior recorded during real–world social interactions, such as those between patient-participants and clinicians, can enable investigation of the neural correlates of arousal and attention in ecologically valid and clinically salient settings.

### Ethical Considerations

Ethical considerations presented by research involving video and audio recording of real-world behavior in a clinical environment include issues of privacy protection, data sharing and publication of findings, and challenges of obtaining informed consent (Berg et al., 2001; Degenholtz et al., 2002). Studies involving such recording affect the privacy of not only participants, but also the visitors, clinicians, and researchers with whom they interact. We believed, and the institutional review board concurred, that with regard to those interacting with participants, this study met the criteria for waiver of informed consent, because obtaining consent was impracticable and the study presented only minimal risks to visitors and others interacting with participants. Instead, a notice was placed on the door of patient rooms to alert anyone entering the room that video and audio recordings would be acquired. Visitors could opt-out by not visiting, or by requesting that their visit not be one of the interactions recorded (perhaps by rescheduling the visit). Clinicians were not able to opt-out of entering and being recorded, as they were required to provide standard care; however, they were informed in advance that the study was being conducted and could raise concerns about their presence and interactions being recorded. These concerns are addressed on a case-by-case basis. (One can imagine, for example, that for reasons of personal safety a clinician might not want her employment location to be made public through future publication/presentation of study findings.) Moreover, the faces of those interacting with participants are to be obscured in all tapes/photos that are either shared or published.

The risks to participant privacy were more substantial, and were simultaneously compounded and mitigated by the clinical environment. In comparison to home environments, inpatient settings afford a lower expectation of privacy, with hospital staff coming and going, rooms often left open to the hallway, and, in some cases, rooms being under video and audio monitoring for reasons of clinical care. Patients generally trade-off their privacy for the prospect of clinical benefit. Nevertheless, the study involved greater reduction in privacy and for reasons that afforded no direct benefit to the participants themselves.

Participants were asked to give informed consent to study participation, including the video and audio recording, collection of physiological data, data sharing, and publication of study findings. Study procedures — putting on, calibrating and wearing the eye-tracking glasses — served to remind participants that their behavior was being recorded. At the end of each recording session, patient-participants were asked to consider the events that happened and explicitly consent to the recording being used for research purposes. In addition, separate consent/release was acquired for use of the video and audio recordings in figures for publications or in presentations. This is especially important because the study took place in a particular clinical setting, and thus for participants who are identifiable in the recordings, publication/presentation of findings would reveal health-related information about them—namely that they were in an Epilepsy Monitoring Unit.

The question of data sharing for recordings that are inherently not de-identifiable is an additional issue to consider. Processed data (annotations with identifiable information removed, for example audio diarization and generic aspects of the computer vision annotations) could likely be shared openly as long as substantial care was taken to assure de-identification. Sharing raw data is a bigger challenge and would require additional layers of consent such as consent procedures used when creating public behavioral databases, though even with this level of protection care must be taken given the potential sensitive nature of the recordings in a clinical environment. Thus, at most, well curated snippets of raw data may be publicly shareable, and sharing of raw data would likely have to be done under IRB approval at both institutions with a data use agreement.

In this study, we sought to study natural real-world social interactions and thus avoided recording doctor-patient interactions or clinical events. For studies that seek to understand doctor-patient interactions or clinical events, these protections and privacy concerns become even more acute and participants should be reminded when acquiring both pre- and post- session consent that the video/audio recordings will include sensitive clinical information.

### Implications for Clinical and Translational Research

Real-world social interactions in an inpatient hospital setting include caregiver–patient interactions (Gert et al., 2021; Girard et al., 2021; Muszynski et al., 2020), which include interactions with neurosurgeons and epileptologists in the case of patients in the EMU. Capturing physiological and behavioral data corresponding to these interactions offers a unique opportunity to understand how clinical decision making in these dyadic interactions is affected by different circumstances based on factors like the severity of clinical issues involved, the presence of family, the patient’s mental health. A deeper understanding of the relationship between patient physiology and behavior that accompanies clinically important interactions has profound implications for clinical practice, patient outcomes and patient satisfaction (Korsch et al., 1968). Lastly, the described workflow can be applied to better understand seizure semiology, which is the keystone for seizure localization and directly related to optimal post-operative results in curative epilepsy surgery.

### Neural Basis of Real-World Behavior

Ecological validity is essential to the investigation of social behavior in the real world. The experimental paradigm we describe here is part of an emerging effort to address this challenge (Babiloni and Astolfi, 2014; Hasson and Honey, 2012; Matusz et al., 2019; Stangl et al., 2021; Topalovic et al., 2020). Laboratory psychology and neuroscience allows for tightly controlled experiments that are crucial for the advancement of knowledge and many aspects of what is discovered in these tightly controlled experiments have external validity (Anderson et al., 1999). However, an ecological approach often yields results that differ from those of laboratory experiments (Gibson, 1979; Holleman et al., 2020; Matusz et al., 2019; Powell and Rosenthal, 2017; Zaki and Ochsner, 2009). For example, recent studies have shown that eye gaze patterns for static faces or even movie faces are very different from those observed during actual face-to-face interactions (Kuhn et al., 2016; Macdonald and Tatler, 2018; Pönkänen et al., 2011; Risko and Kingstone, 2015; Risko et al., 2012, 2016) and real world settings have been shown to activate broader brain networks than do artificial conditions (Camerer and Mobbs, 2017; Hasson and Honey, 2012; Nili et al., 2010). Moreover, the “naturalistic intensity” (Camerer and Mobbs, 2017) of an interaction with one’s loved ones or a doctor or a threatening stranger is a key element of real-world experience that cannot be fully captured in a laboratory. Basic aspects of the organization of the “social brain” (Wang and Olson, 2018) are unlikely to change in real-world environments, for example regions of the brain that show face selectivity in the lab (Boring et al., 2021; Kanwisher, 2000; Tsao and Livingstone, 2008) remain face selective in natural conditions (Fig. 7), as expected given that disruption to these regions cause real-world face processing abnormalities (Barton et al., 2002; Parvizi et al., 2012; Zhang et al., 2015). However, important aspects of how these regions code and process social information are likely to reflect real-world processes that differ from the laboratory environment. At a minimum, it is important to validate laboratory findings in real world settings to determine the generalizability of models derived from controlled experiments (Anderson et al., 1999).

The complexity of studies in the real-world is that there is enormous uncontrolled variability in natural environments. However, modern computational studies, such as those in artificial intelligence and computer vision, show that real-world variability can be well-modeled with sufficient data. Our paradigm is designed to enable real-world neuroscience by facilitating the collection and processing of large datasets combining behavior, physiology, and neural recordings that can be analyzed using modern computational techniques to test hypotheses about social behavior and its neural bases in natural environments.

The movement towards studying the neural basis of real-world behavior has also been seen in recent studies with non-human subjects, enabled by the potential of telemetric recordings that allow for neural activity to be recorded during natural behavior (Fernandez-Leon et al., 2015; Mavoori et al., 2005; Roy and Wang, 2012). Parallel studies of natural neuroscience in non-human primates has the potential to allow for a deeper understanding of the cellular-to-systems mechanisms for basic pan-specific aspects of social behavior and cognition. Advances in computer vision provide the opportunity to annotate nonhuman animal behavior and in relation to details of a natural environment (Mathis et al., 2018) just as they do in human studies. Recent work has also demonstrated that restraint free, real-world eye tracking is also possible in non-human primates (Hopper et al., 2020; Ryan et al., 2019). Thus, the approach described in this work could be adapted to parallel studies in non-human primates, leveraging the higher resolution methods that are possible to use in nonhuman primates, to allow a cellular-to-systems understanding of the neural basis of real-world cognition and perception.

## Conclusion

We view the approach outlined above as part of an ongoing paradigm shift in approach towards studying real-world behavior and cognition and their neural underpinnings. Real-world ”naturalistic intensity” and ecological validity is particularly important for studying social interactions and their neural correlates. Our current methodology augments eye-tracking and behavioral monitoring in experimental recording sessions in the EMU with neurophysiological monitoring. Extending behavioral monitoring to unscripted and more real-world contexts can enable the collection of multi-modal data sets that are large enough for cutting edge machine learning techniques like deep learning to be pressed into service to learn relationships between behavior and physiology. Combined behavioral and physiological data can be used both for studying basic cognitive phenomenon and can also be used to find markers that are predictive for clinically significant events like seizures, cardiac events, respiratory events and others.

## Acknowledgements

We thank the patients for participating in the iEEG experiments; the UPMC Epilepsy Center monitoring unit staff, physicians and administration, particularly Cheryl Plummer, Maggen Millen, and Dale Crawford, for their assistance and cooperation with our research; Nicole Silverling and Taylor Gautreaux for their help with annotations of the egocentric video; and Katherine Lindsay for her help with audio annotations.

## Open Practices Statement

Data are available upon reasonable request and IRB approval.

Custom python scripts to parse eye-tracking traces, construct an eye-tracking event log and reconcile gaze location with computer vision annotations (bounding boxes) to identify each fixation as a face or non-face fixation are available (with sample data and output) at https://github.com/arishalreja/lcnd_eye_tracking_real_world_vision.

YOLO v3, the computer vision model used for object detection is available at https://pjreddie.com/darknet/yolo/.

Open Face, the facial behavior analysis toolkit for extraction of facial landmarks, head orientation and position, eye gaze direction and facial expression, is available at https://github.com/TadasBaltrusaitis/OpenFace.

Experiments were not preregistered.

